# Overlapping neural processes for stopping and economic choice in orbitofrontal cortex

**DOI:** 10.1101/304709

**Authors:** Pragathi Priyadharsini Balasubramani, Benjamin Y. Hayden

## Abstract

Economic choice and stopping are not traditionally treated as related phenomena. However, we were motivated by foraging models of economic choice to hypothesize that they may reflect similar neural processes occurring in overlapping brain circuits. We recorded neuronal activity in orbitofrontal cortex (OFC), while macaques performed a stop signal task interleaved with a structurally matched economic choice task. Decoding analyses show that OFC ensembles predict successful versus failed stopping both before the trial and immediately after the stop signal, even after controlling for value predictions. These responses indicate that OFC contributes both proactively and reactively to stopping. Moreover, OFC neurons’ engagement in one task positively predicted their engagement in the other. Finally, firing patterns that distinguished low from high value offers in the economic task distinguished failed and successful trials in the stopping task. These results endorse the idea that economic choice and inhibition may be subject to theoretical unification.

## INTRODUCTION

Stopping (sometimes referred to as inhibition) and economic choice are two major brain functions that have historically been studied independently. Nonetheless, there is some reason to think that they may spring from shared processes. For example, several psychiatric conditions, including depression and addiction, impair both processes, and greater impairment of both is associated with greater disease progression (Iacono et al., 2008; Nestler et al., 2002; Volkow et al., 2011). Second, both are closely associated with, and empirically linked to, the broader concept of self-control (Berkman et al., 2016; Hayden, 2018; Inzlicht et al., 2014; Shenhav, 2017). Third, both tend to activate a similar set of brain regions, including the pre-motor cortex, the ventrolateral prefrontal cortex, basal ganglia, and the thalamus (Aron, 2007; Aron and Poldrack, 2006; Cisek, 2012; Cisek and Kalaska, 2010; Hampshire and Sharp, 2015; Sakagami and Pan, 2007; Schall et al., 2002).

The idea that stopping and choice may have a deeper relationship is motivated by certain foraging-inspired approaches to economic choice (Cisek, 2012; Cisek and Pastor-Bernier, 2014; Hayden, 2018; Hayden and Moreno-Bote, 2018; Kacelnik et al., 2011; Krajbich et al., 2010; Stephens and Krebs, 1986). A core tenet of these approaches is that the brain’s decision-making systems are evolved to make accept-reject decisions (Kacelnik et al., 2011; Ojeda et al., 2018; Pirrone et al., 2017; Shapiro et al., 2008; Vasconcelos et al., 2010). Even ostensibly binary economic choices, in this view, reflect a pair of (potentially interacting) accept-reject choices. Each accept-reject decision, in turn, involves choosing whether to pursue an option or refrain from pursuit. *Accepting* involves selecting the attended or activated option, or, more abstractly, performing the afforded action (Cisek and Kalaska, 2010; Cisek and Pastor-Bernier, 2014; Hayden and Moreno-Bote, 2018). *Rejecting* involves countermanding the afforded action. A classic binary economic choice, then, may be seen as two related decisions about whether to go or stop choosing the attended option or the afforded action (Hayden, 2018).

This way of looking at choice is consistent with some recent studies that suggest that binary choice involves a serial, not parallel, consideration of options (Krajbich et al., 2010; Rich and Wallis, 2016; Strait et al., 2014; reviewed in Hayden and Moreno-Bote, 2018). These studies and others indicate that attention is largely limited to a single option, which is evaluated, often relative to the other one (Lim et al., 2011; Rich et al., 2017; Rudebeck and Murray, 2014; Strait et al., 2014 and 2015; Xie et al., 2018). Choice, then, presumably occurs relative to a single option that is evaluated relative to a background value, which includes the value of choosing the other option (Shapiro et al., 2008; Vasconcelos et al., 2010). However, this work does not directly tie economic choice and reward processing to stopping processes.

Here we sought to test the overlap hypothesis by comparing neuronal activity in an economic choice task with that observed in a stopping task. We focused on the orbitofrontal cortex (OFC). The centrality of OFC in economic choice is largely undisputed, although its specific role remains to be determined (Padoa-Schioppa, 2011; Rich et al., 2017; Rudebeck and Murray, 2014; Schoenbaum et al., 2009; Wallis, 2007; Wilson et al., 2014). It is clear, nonetheless, that activity of Area 13 of OFC correlates with the values of offers and of chosen options, and is likely to be critical for value comparison as well (Padoa-Schioppa, 2013; Padoa-Schioppa and Assad, 2006; Raghuraman and Padoa-Schioppa, 2014). In contrast to its clear role in choice, the contribution of the OFC to stopping remains disputed. On one hand, a good deal of work argues against a direct inhibitory role for the OFC (Chudasama et al., 2006; Ghods-Sharifi et al., 2008; Rudebeck and Murray, 2014; Schoenbaum et al., 2003; Stalnaker et al., 2015). However, multiple studies give the OFC at least some role in inhibition (Bryden and Roesch, 2015; Chikazoe et al., 2009; Dias et al., 1996; Eagle et al., 2007; Horn et al., 2003; Majid et al., 2013; Mishkin, 1964; Roberts and Wallis, 2000). One reason for the continued debate about the role of OFC in choice the lack of direct evidence from the unit activity in this region in stopping tasks (but see: Bryden and Roesch, 2015).

We hypothesized that OFC participates in both choice and stopping decisions in similar ways, that is, by computing executive signals that promote (or fail to promote) particular strategies. To test this hypothesis, we examined responses of OFC neural populations recorded in two interleaved tasks, a stop signal task and an economic choice task. The tasks were designed to have structures as similar as practically possible. We were particularly interested in the questions of (1) whether and how the function of this economic region includes stopping, and (2) whether neural response patterns related to stopping correspond with patterns related to value.

## RESULTS

### Behavior in the stop signal task and economic choice task

Subjects performed a ***standard stop signal task*** (similar but not identical to the one used by Hanes and Schall (1995; **Figures 1A** and **1B** and **Methods**). On each trial, following a central fixation, subjects saw an eccentric target (*go signal*) that, if fixated, provided a juice reward. On a subset of trials (33%, called stop trials), a second signal (*stop signal*) appeared at fixation and countermanded the previously instructed saccade. Successful stopping trials were rewarded. Failed trials (trials in which a saccade was made despite a stop signal) were not. Median reaction time to the target in go trials was 0.435 sec and 0.272 sec in subject J and subject T, respectively (**Figures 1C** and **1D**). Average trial length including feedback time for subjects J and T were 3.66 s and 2.62 s, with the mean feedback start time as 1.78 s and 1.49 s, respectively. Both subjects showed typical behavior in this task; their performance in stop trials varied as a function of time of presentation of stop signals relative to that of go signal (**Figures 1E** and **1F**).

**Figure 1.**
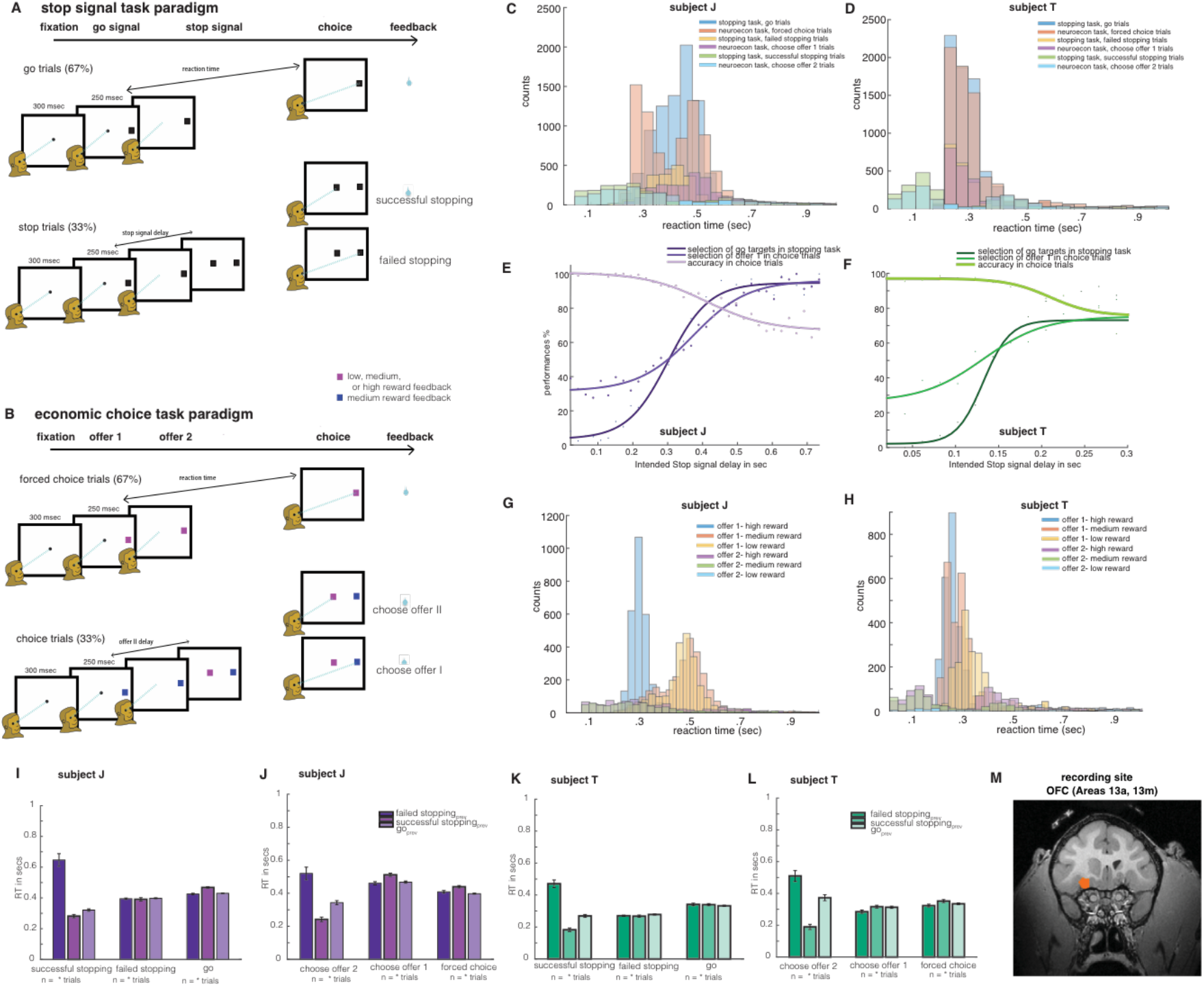
Task, anatomy, and subject behavior: task framework **(A)** stop signal task **(B)** economic choice task. Behavioral results for subject J are presented in panels (C, E, G, I, J) and for subject T in (D, F, H, K, L) (**C, D**) reaction time distributions for various trial conditions of stop signal task (E, F) Inhibition function and accuracy of choices varied as a function of SSDs (**G, H**) reaction time distributions for various trial conditions of economic choice task. Previous trial had effects in reaction time behavior in (**I, K**) stop signal task and (**J, L**) economic choice task. Error bars represent SEM, and * denotes t-test significance with p < 0.05. (M) recording site – Area 13 of OFC (scan from subject J shown).

The delay between the go signal and the stop signal is called the stop signal delay (SSD) and it varied randomly across trials. We estimated the SSD that leads to approximately 50% successful stopping (SSD-50) because it can help in computing the stop signal reaction time (SSRT, Logan, 1994; Logan and Cowan, 1984; Verbruggen and Logan, 2008). The SSD-50 was 0.277 sec for subject J and 0.131 sec for subject T. SSRT computed for SSD-50 was 0.158 sec for subject J, and 0.141 sec for subject T. These values are typical of rhesus macaques in these tasks (e.g. Hanes and Schall, 1995; Ito et al., 2003).

We randomly interleaved stop signal trials with trials from an ***economic choice task***. This task was designed to have a similar structure to the stop signal task. Specifically, forced choice trials were equivalent to go trials and choice trials were equivalent to stop trials (see **Methods** for details). In the economic choice task, the offers were associated with low (yellow), medium (blue), or high (magenta) reward value (**Figure 1B**). The subjects chose either offer 1 (which appeared at the periphery, similar to go signal) or, when it occurred, could choose the later-appearing offer 2 (which appeared at the center, similar to stop signal). The delay for offer 2 was fixed and defined by the measured stop signal delay computed from the stopping task. For simplicity, we will use the terms accept and reject trials to refer to those in which the subject chose offer 1 and offer 2, respectively.

As anticipated, the two tasks showed similar behavior results. Median reaction time in forced choice trials was 0.41 sec and 0.27 sec in subject J and subject T, respectively (**Figures 1G** and **1H**). The reaction time medians for choice trials in the presence of offer 2 were lower that that in forced choice trials (**Supplemental results-A**). On average, the total length of trials including the feedback time for subjects J and T were 3.86 s and 2.88 s, with the mean feedback start time as 1.88 s and 1.70 s, respectively. Choice accuracy varied as a function of SSD in both subjects (**Figures 1E** and **1F; Supplemental results-A**) Both subjects showed similar behavioral effects in the current trial of each task, as a function of previous trial conditions (**Figures 1I** and **1J** for subject J, **Figures 1K** and **1L** for subject T; refer to **supplementary results A** for behavioral effects).

### Overlapping sets of neurons participate in the two tasks

We recorded responses of 96 neurons (52 in subject J and 44 in subject T) in Area 13 of the OFC (**Figure 1M**). The number of neurons to be collected was determined *a priori* based on exploratory analyses of previously collected datasets and was not adjusted during recording based on analyses performed mid-experiment. Note that while this number is smaller than in some other studies, is it sufficient to detected the effects we are interested in here.

We focused our analyses on four key time periods of the trial: (1) the 300 ms epoch *before the trial* started (*pre-go signal epoch*), which corresponds to fixation time before the appearance of any stimulus targets; signal differences here presumably reflect proactive control (Stuphorn and Emeric, 2012). (2) The variable time after the *stop* signal and before the reaction time period (*post-stop signal epoch*). The post-stop epoch is important because it is when inhibition generated in response to countermanding commands would presumably occur, and has therefore been the focus of many studies of stopping (Logan et al., 2015; Schall, 2001; Schall et al., 2002). It corresponds to the time during which reactive control occurs (Stuphorn and Emeric, 2012). (3) The variable time after the go signal and before the reaction time period (*post-go signal epoch*). The equivalents of (2) and (3) in choice task are the variable times after *offer 2*, and *offer 1*, respectively before the reaction time of a trial; these epochs denote the task related activity in general; (4) the variable time between the beginning and end of feedback reward (*feedback epoch*).

Neurons had diverse tuning profiles with near balance of sign. In stopping task, 44.79% of cells showed positive task tuning (response during task versus baseline), and 53.12% showed negative tuning, and the rest weren’t modulated. In the economic choice task, 45.83% of cells showed positive task tuning, and 51.04% showed negative tuning, and the rest weren’t modulated. We examined the relationship between simple response patterns in the two tasks on a cell-by-cell basis. We found similar neuronal activity when comparing task related activity against baseline across tasks (which we call *task-related tuning*). Regression weights (task-related tuning coefficient) in one task positively predicted the weights in the other task (Pearson correlation between tuning coefficients, r = 0.78, p < 0.001, Figure 2A). (Note that these data come from separate sets of trials, so there is no overlap in data used to estimate the two sets of tuning coefficients.) Moreover, absolute response patterns (that is, unsigned regression weights) were also positively correlated across the two tasks (0.74, p < 0.001), indicating that it was the same set of neurons involved in the two tasks, rather than distinct sets (for details on using this method to interpret relationships between regression coefficients, see Blanchard et al., 2015a).

**Figure 2.**
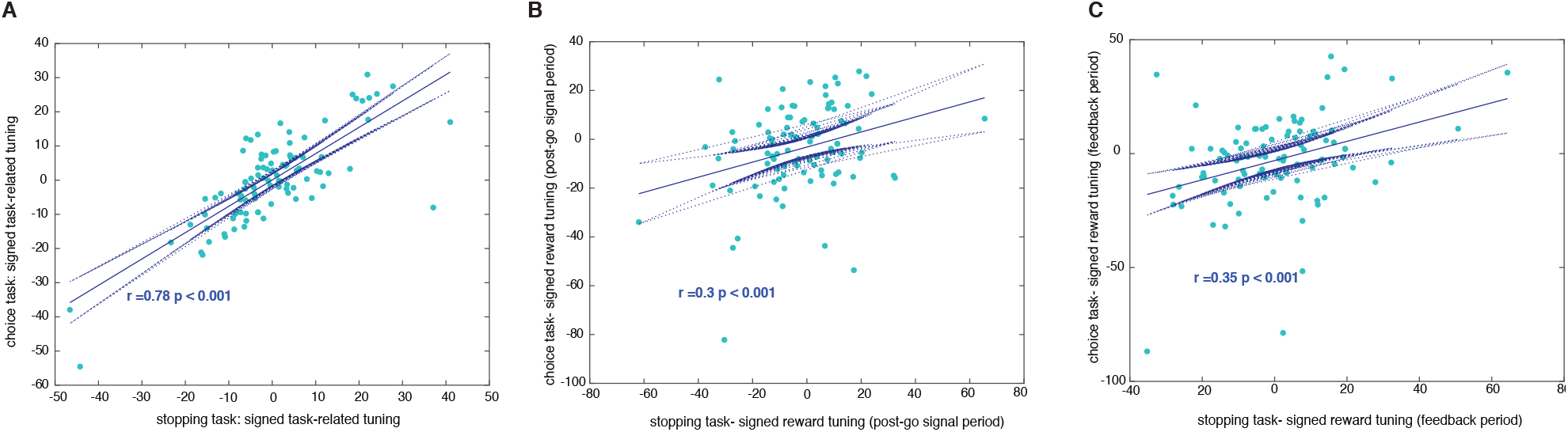
overlapping sets of neurons across stopping and choice tasks: Correlations between signed **(A)** task-related tuning coefficients, reward tuning coefficients in **(B)** post-go signal epoch **(C)** feedback epoch of stopping and economic choice tasks.

Next, we compared the coding of rewards in both tasks. For the stop signal task, we looked at the differential coding of no-rewards during failed stopping versus rewards in successful stopping; for the economic task, we looked at differential coding of varied rewards associated with offers. Tuning coefficients for reward values were positively correlated between both the tasks during the post-go signal period (between signed coefficients, r = 0.3, p < 0.001, **Figure 2B**) and during the feedback epoch (between signed coefficients, r = 0.35, p < 0.001, **Figure 2C**). The positive correlation demonstrates that the rewards are encoded in similar way across tasks. Moreover, the unsigned correlation coeffcients were also correlated in both epochs (r = 0.27, p = 0.01 and r = 0.28, p = 0.01, respectively). This correlation indicates that coding of the two types of value was handled by the same subset of neurons across the two tasks.

### Selectivity for stopping in single neurons

The role of OFC in economic decisions is well established, but its role in stopping is not. Thus, a demonstration of functional overlap involves showing that it plays a role in stopping. Because of its relevance to the foraging hypothesis (see **Introduction**), we focused here on the determination of successful from failed stopping. The two time periods of significant interest to this hypothesis include: 1) The post-stop signal period, and 2) the pre-go signal time period.

Responses of example neurons are illustrated in **Figure 3**. In neuron J19, firing rates following the go signal but before the SSRT were lower on successfully inhibited trials (1.8 spikes/sec) than on failed stopping trials (4.1 spikes/sec, Wilcoxon rank test, ranksum = 1480, p < 0.05, n = 567 trials, **Figure 3A**). Note that there is a larger and more prominent modulation in firing rate later in the trial. Given its timing, this modulation likely relates to outcome monitoring and is too late to influence stopping. Another example neuron, T25, showed distinct patterns for successful and failed stopping trials even 500 msec before the beginning of the trial (ranksum = 2080, p < 0.05, n = 579 trials, **Figure 3B**).

**Figure 3.**
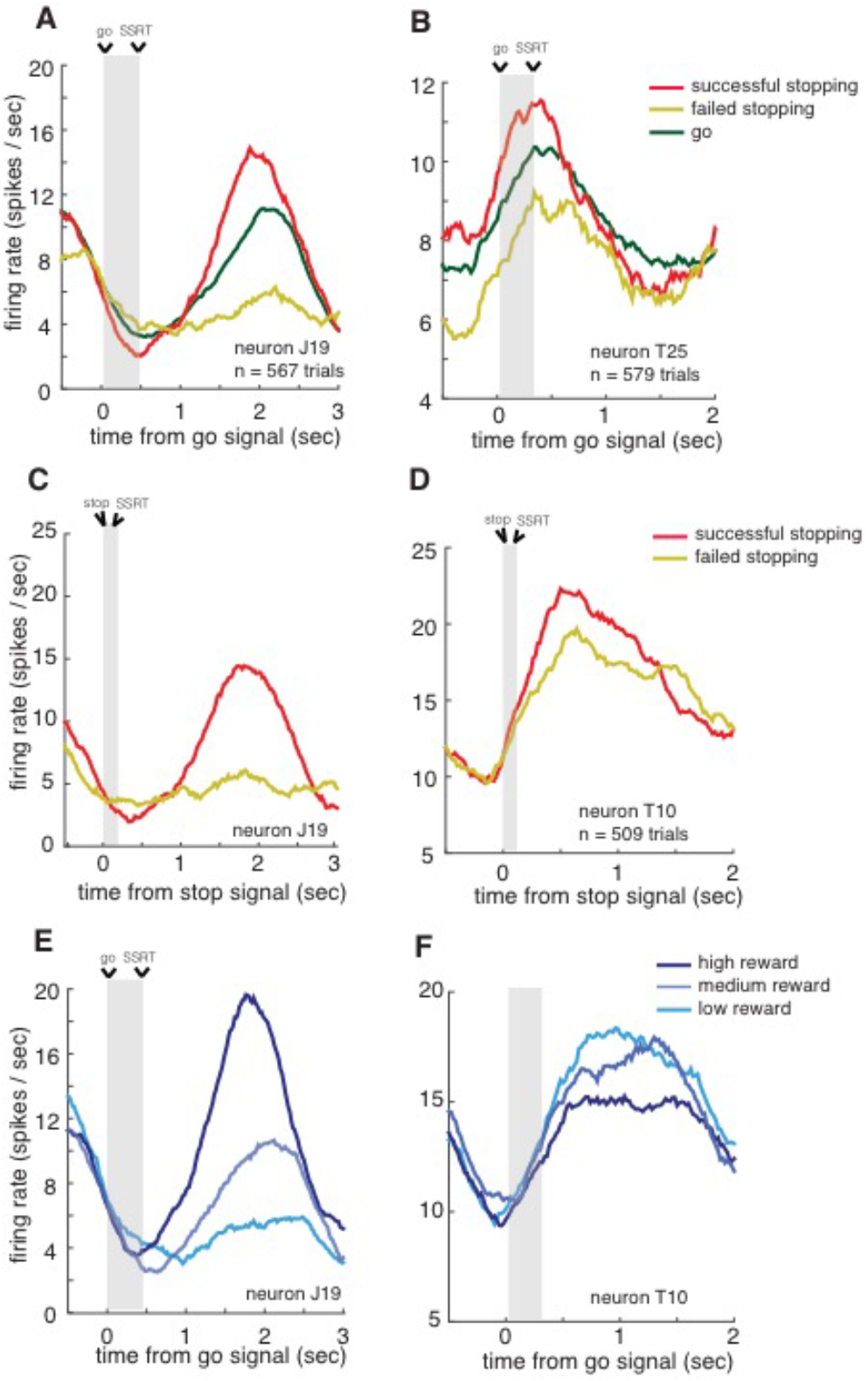
selectivity for stopping in sample neurons: Activity of example neurons during successful stopping, failed stopping and go trials are illustrated with respect to go signal in panels (**A, B, E, F**) and stop signal in (**C, D**) presentation time. Time from start of the go (stop) signal to SSRT is shaded all panels. Neuron in panel A shows significant difference in firing rates of successful and failed stopping trials before SSRT. Neuron in panel B shows difference even before the beginning of trial. Neuron in panel **C** is the same as panel A, shows significant difference in firing rates of inhibition trials before stopping response time. Likewise, neuron in panel D shows difference around few msecs after SSRT. (**E, F**) Activity of example neurons in economic choice task: neuron in panel **E** is the same as panels **A** and **C**. Its reward related activity after SSRT in the choice task parallels to that of stop signal task, and is positively correlated to the value. Neuron in panel F shows the opposite trend, and is negatively correlated to the reward value.

The responses shown in **Figure 3C** and **3D** are aligned to stop signal (time zero). Figure 3C illustrates the activity of the same neuron shown in **Figure 3A**; its response pattern showed significant differences between successful stopping trials (1.8 spikes / sec) and failed stopping trials (4.4 spikes / sec) that begin after the presentation of stop signal but before SSRT (ranksum = 1340, p < 0.05). Finally, neuron T10 (**Figure 3D**) fired more vigorously on successful than on failed stopping trials at around 100 msec after the SSRT (ranksum = 2229, p < 0.05). The results also show that in OFC, the codes for go and stop trials do not show simple and opposite activities for a saccade. This contrasts with the activities of movement and fixation neurons in FEF, where inhibition is driven by a rapid rise in firing rates of a specific subpopulation of neurons—fixation neurons – that gate the activity of another subpopulation—movement neurons (Hanes and Schall, 1995; Logan et al., 2015; Schall, 1991). Therefore, OFC neurons are rather complex for a race model (Logan et al., 2015) to compute their stopping pattern.

To test for the possibility that our putative inhibition signals were just reward correlates, we took advantage of trials collected from the economic choice task. The data from this task allowed us to assess each neuron’s tuning function for anticipated rewards. Responses to different reward amounts by two example neurons are shown in **Figures 3E** and **3F**. We found tuning for anticipated reward values in the firing activity during the reward feedback time period. For example, we observed a significant positive correlation between reward amount and firing rate in neuron J19 (ϱ = 0.3138, p < 0.001, **Figure 3E**), this neuron shows similar reward related activity across tasks (**Figure 3A, 3C**). We also show another neuron with a significant negative correlation to rewards (neuron T10, ϱ = – 0.143, p = 0.04, **Figure 3F**). Example individual neurons suggest the presence of diverse stopping and reward related neuronal codes at OFC. Furthermore, simple population analyses don’t inform about stopping patterns at OFC (refer to **supplementary results-B**).

### Ensemble patterns distinguish successful from failed stopping

To compare whether similar ensemble patterns of activity predicted behavior in the two tasks, we examined neural network decoders (classifiers). (Note that while we used normalized data, see **Methods**, the normalization procedure did not alter our results, see **Supplementary results-C**). We first trained decoders to analyze differences in population activation patterns between successful and failed stopping in stop signal task; We trained classifiers using two key epochs— *post-stop signal epoch* and *pre-go signal epoch*. Testing of decoders for significant patterns differentiating successful and failed stopping used 100 msec moving boxcars stepping in 10 msec intervals as input to the classifier.

Because of the possibility of false positives, we were especially interested in periods in which a decoder had several positive effects in adjacent bins (see **Methods**). The post-stop signal decoder was able to classify success versus failure of inhibition significantly in a series of 9 consecutive bins spanning 40 to 120 msec after the stop signal (these times indicate the starts of the 100 msec boxcars; see **Methods** on procedures to determine statistical significance of a boxcar using chi-square statistics). The corresponding numbers for individual subjects were 40 to 140 msec in subject J and 40 to 220 msec in subject T (**supplementary results-D**). The post-stop signal pattern series are unlikely to occur by chance (p < 0.001 in all cases, see **Methods** for specific use of chi-square tests to quantify significance of consecutive bins). Since each bin spans 100 msec, the central point of the first element in this series provides a rough estimate of the latency of the effect. That number was t = 90 msec for both subjects.

Notably, the central point of the first bin of the series to reach significance in both subjects occurred before the stop signal reaction time of either subjects (the SSRTs were 140 msec for subject J and 120 msec, also see **supplementary results-D; Figure 4**). We call the central point the cancellation time; it measures the center point latency of first statistically significant difference between successful and failed stopping trials for the ensemble of neurons. The cancellation time is 90 msec for both subjects. The cancellation time preceded the average stopping response (SSRT) by 50 msec in subject J, and by 30 msec in subject T, suggesting OFC’s stopping-related patterns precede the stopping response (see Discussion, and **supplementary results-D**).

**Figure 4.**
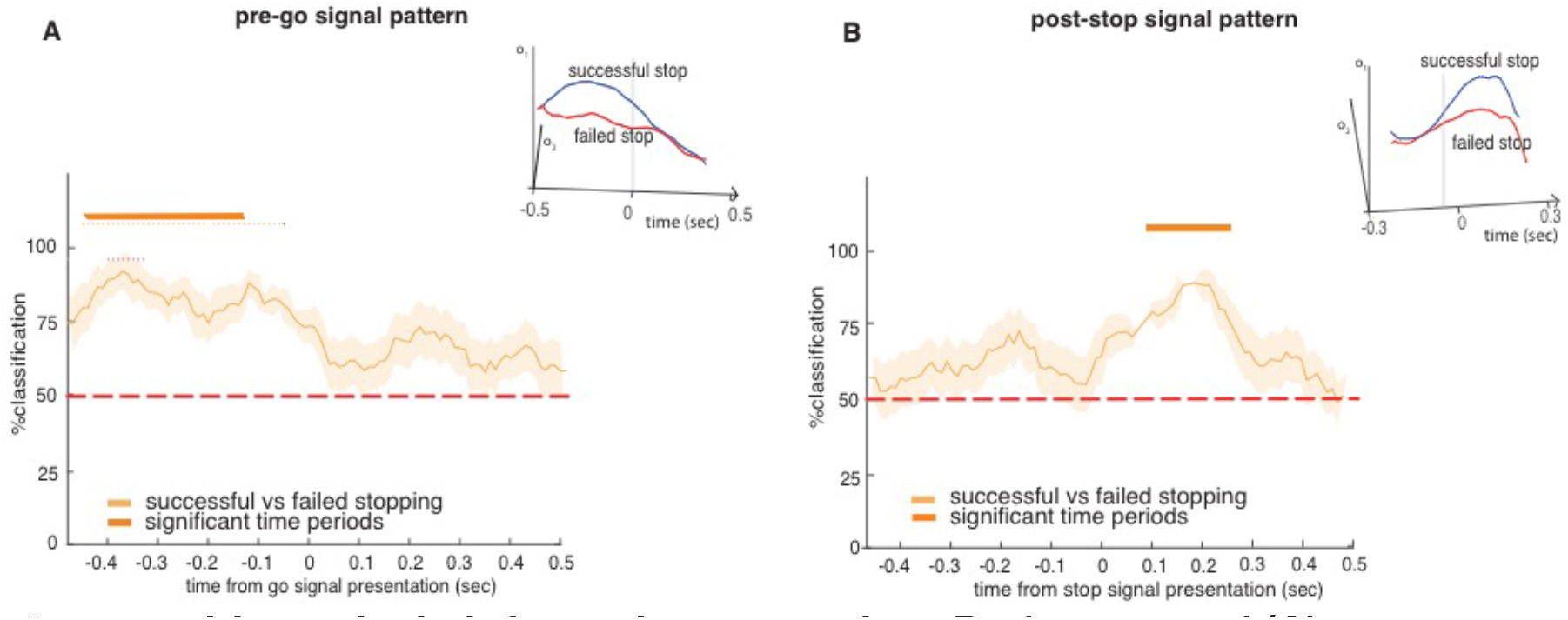
ensemble analysis inform about stopping: Performance of **(A)** pre-go signal **(B)** post-stop decoders to distinguish successful versus failed stopping pattern. Insets in both panels illustrate sample projections of decoder’s output responses (Y-and Z-axes indicate the values of o_1_ and o_2_, for successful and failed stopping trials, respectively). Error bars indicate SEM, so non-overlap with the chance bar (horizontal dashed red line) is not sufficient to indicate statistical significance). Time points underneath through thick horizontal orange bars (and black tick marks) denote start time of 100 msec boxcars having classification accuracy significantly above chance (i.e. 50%) (chi-square test, p < 0.05).

We then examined the response differences of the pre-go signal decoder. We observed a significant pattern difference between successful and failed stopping trials (p < 0.001) in 36 number of boxcars, extending from 470 to 120 msec before the go signal (see **Methods** for details on chi-square statistics). For subject J, significant decoding was observed in 35 number of boxcars during the time periods 460 to 120 msec; for subject T it was 23 number of boxcars from 420 to 200 msec. These results indicate that the upcoming success or failure of inhibition is decodable from OFC patterns even before the start of the trial (**Figure 4**, also see **supplementary results-D**). The pre-go signal pattern series are unlikely to occur by chance (p < 0.001 in all cases, see **Methods** for specific use of chi-square tests to quantify significance of consecutive bins). Our results do not tell us why this correlation exists, although one may infer that it reflects some internal state that drives successful versus failed inhibition. Thus is it is a likely correlate of proactive control. Overall, these results implicate Area 13 of OFC in the process of regulating stopping decisions.

### The post-stop and pre-go decoders are statistically orthogonal

We next examined how the pre-go and post-stop decoders related to each other. We did so by comparing their weight vectors. We found a very low similarity between them (Pearson correlation coefficient, r = 0.0008± 0.0086), suggesting the two epoch patterns may be nearly statistically orthogonal and hence independent. A different possibility is that this low correlation may be an artifact of noise. To test this second possibility, we next performed a cross-validation procedure to estimate the maximum range of measured cross-correlation values had the variables been fully correlated given our noise properties. Cross correlation coefficient obtained between converged weight values of pre-go and post-stop signal decoders (theoretical maximum), *‘r_xmaX_* is 0.9058 ± 0.0878; the value fell outside the central 98% of cross-validated data (and is significant at p <= 0.01; 100 randomizations, average of *r_x-max_* from the randomized sets = 25.84 ± 0.82), substantiating statistical independency between pre-go and post-stop signal decoder weight patterns.

Moreover, when the decoding performances for two decoders were compared, they were significantly different during the time periods after the stop signal (t-stat = 6.0491, p = 0.003). Similarly, they were significantly different even before the beginning of trial (t-test, t-stat = 8.8874, p < 0.001). The above results suggest that the two decoders that predict successful versus failed inhibition are statistically orthogonal and thus dissimilar. More broadly, these results suggest that the computations associated with ostensible proactive and reactive control of stopping by OFC are distinct.

### OFC encoding of stopping is not a by-product of value coding

The reward-encoding role of OFC is a hallmark of its function (Padoa-Schioppa, 2011; Rushworth et al., 2011; Schoenbaum et al., 2009; Schoenbaum et al., 2011; Schultz, 2000; Wallis, 2007). We therefore wondered whether the stopping-related activity that we observed might by an artifact of its reward roles. For example, it may be that there is some undetectable natural variation in the relative subjective value of the reward offered for correct performance. On trials in which the reward happened to have a slightly lower value, the subject would be less motivated to perform correctly; this fluctuation would then introduce a correlation between firing rates and successful stopping (see, for example, Sugrue et al., 2005, for a similar argument about LIP neurons).

If the stopping-related signals were a direct consequence of reward encoding for every neuron, we would see a positive correlation between coding patterns for rewards and stopping. We computed a *reward index* for all neurons by regressing their responses at feedback epoch against the outcomes themselves. We computed a stopping index for all neurons by subtracting on their firing rate during successful and failed stopping before SSRT (see **Methods**). We found no correlations between these indices in the post-stop signal time period (Pearson correlation, ϱ =0.09, p=0.4, **Figure 5**). Nor did we find such correlations in pre-go signal time period (ϱ =-0.02, p=0.82, **Figure 5**).

**Figure 5.**
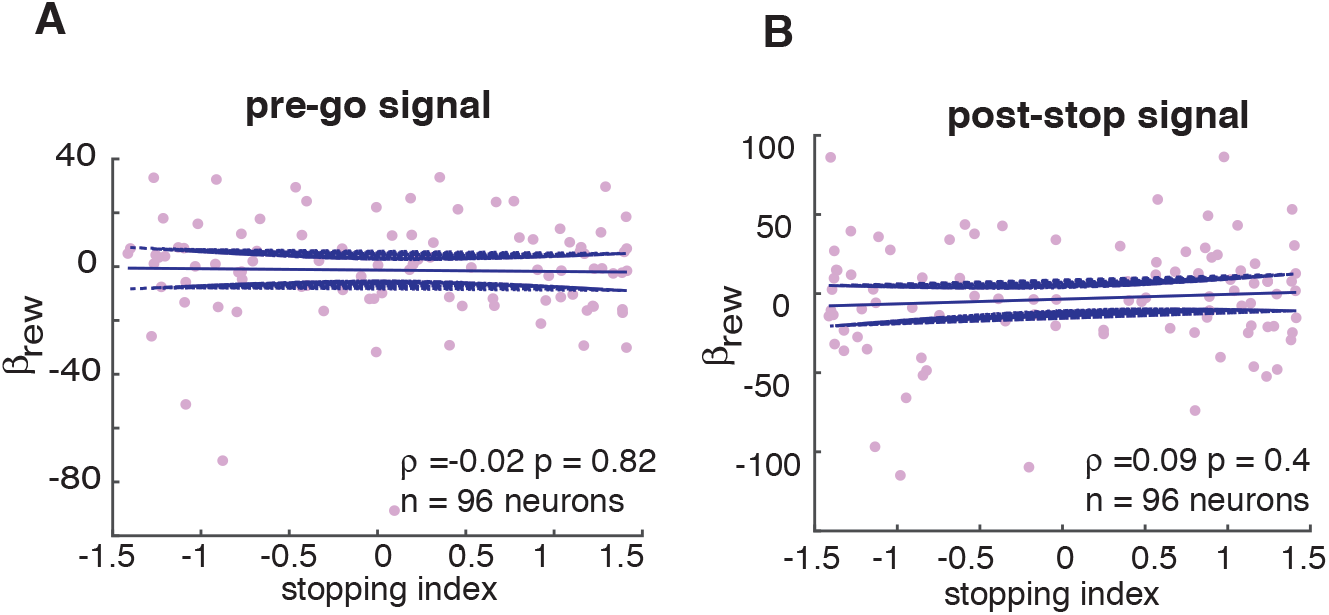
unrelated reward and stopping codes: Correlations between stopping and reward indices show no significant effect during 100 msec in **(A)** pre-go signal and **(B)** post-stop signal time period.

This lack of correlation at the neural level may be a sign that the reward code and the stopping code are different. It may also, in theory, be due to lack of sufficient data to detect a significant effect. To test this idea, we performed a cross-validation analysis (See **Methods**). Specifically, we reasoned that if insufficient data were the problem then a within sample correlation would also produce no significant correlation. A positive coefficient resulting from a within sample correlation, using randomly sampled half-sized subsets, then, would indicate that our data have sufficient power to detect a significant effect (Blanchard et al., 2015a). We thus tested whether the correlation coefficient for stopping and reward indices fell below the bottom 5 percentile of the coefficients obtained for within-group correlations. Indeed, the coefficient fell below 1st percentile of that obtained for 100 randomizations in cross validation analysis. **Figure 5** show nocorrelations between stopping and reward indices with p <= 0.01.

### Overlapping functional ensemble codes for stopping and for value

We next examined the relationship between patterns that distinguished successful versus failed stopping of the stop signal task, and different reward values of the economic choice task. One potential confound in such an analysis is that OFC may encode action, and similar effects may reflect their shared actions (Feierstein et al., 2006; Grattan and Glimcher, 2014; Roesch et al., 2006; Yoo et al., 2018; Strait et al., 2016). To deal with this problem, for the analysis of economic choice task data, we used only trials in which the subject accepted the offer. Thus, action was the same – a saccade – in all cases. Below, we present results on training using two key epochs representing the trial, its task-related activity informing (1) post-go signal epoch, and the (2) the feedback epoch.

We first trained a network with successful versus failed stopping trials, and tested them with high versus low value-accept trial (see **Methods**). Therefore, this decoder network looks for stopping task related patterns in the economic choice trials. When trained with post-go signal epoch, decoder differentiating successful versus failed stopping patterns showed significant similarities to time periods −140 to −60 msec, 420 to 570 msec, from the presentation of offer 1 for patterns differentiating low versus high rewarding accept trials (**Figure 6**). We also trained a reversed-network with reversed training and testing sets: trained with patterns distinguishing high versus low value-accept trials, and tested with patterns distinguishing failed and successful stopping trials. The significance of the reversed network, in contrast with the earlier, looks for the presence of economic choice task related patterns in the stopping task trials. Similar to the earlier, in reversed-network, training with patterns differentiating low versus high rewarding accept trials of choice task, in the post-go signal epoch, showed significant similarity to time periods around −230 to −70 msec, 420 to 720 msec from the presentation of go signal for patterns differentiating failed versus successful stoppings, respectively (**Figure 6**). This shows the presence of overlapping patterns between tasks, the neural patterns relating to the decision process of one particular task-type can be tracked from many time periods (pre trial onset and during the later parts of the trial) of the other task-type.

**Figure 6.**
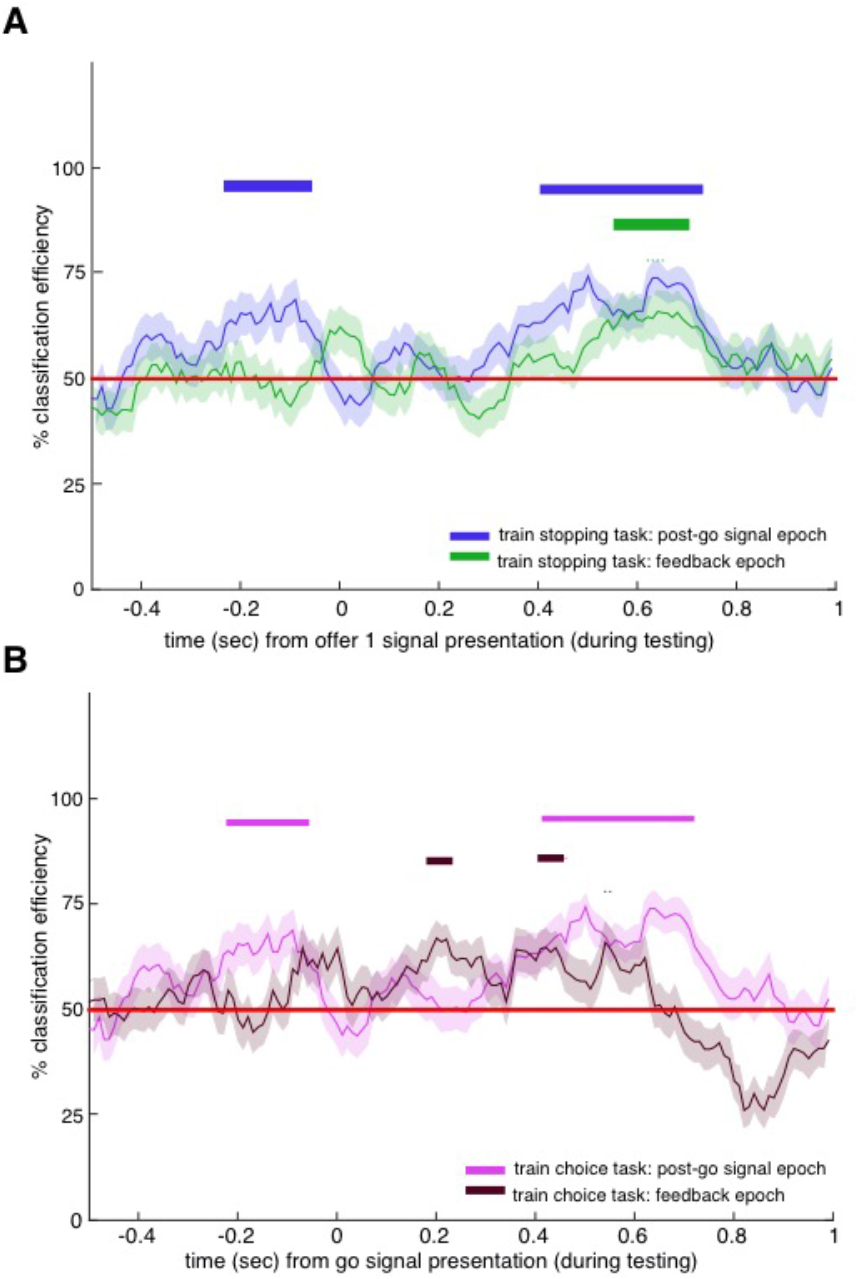
overlapping functional ensemble codes for stopping and choice: Performance of decoders trained on **(A)** stopping task **(B)** choice task on poststop signal epoch (blue, magenta) and feedback (green, brown) epochs. Error bars represent SEM. Time points underneath the thick horizontal highlighted bars (blue, green, magenta, brown) of each result, as labelled in the legend, denote the start time of 100 msec boxcars having percent accuracies of classification above chance of 50% (chi-square test, p < 0.05). Red line indicates chance (50%) performance for both decoders.

Then, we trained with feedback epoch of successful versus failed stopping trials; their patterns showed similarity to time periods 570 to 690 msec after the presentation of offer 1 of high versus low rewarding accept trials (**Figure 6**). In the reversed-network, with characteristics described as the previous, training using high versus low rewarding accept trials of choice task showed significant similarities to time periods around 190-220 msec and 400-440 msec of failed versus successful stopping patterns, respectively (**Figure 6**). These results show the similarities of feedback related ensembles in both tasks.

Reward promotes choice of the option it is associated with. The results show that the patterns used to encode stopping in the stop signal task would correspond to patterns associated with value encoding in the economic choice task (Hayden, 2018). Particularly, we found that task and feedback epoch related patterns of one task can be tracked from the other, in time periods both before and during the trial, in a non-continuous and dispersed manner. Similarities between neural patterns of both tasks may promote the idea of shared strategies between stopping and choice decisions. We hypothesize that reward related patterns could act as motivational forces when present in stopping decisions. On the other hand, presence of stopping related patterns in choice decisions could facilitate internal action strategies guiding the choice.

## DISCUSSION

We examined responses of neurons in Area 13 of the OFC in two tasks, one an implementation of a classic stopping task and the other a simple economic choice task with similar structure. Although the economic role of this region in choice is well established, its role in stopping is not. Our finding that OFC ensembles predict stopping both before the trial and immediately before the stopping response demonstrate that this region does participate in regulation of stopping. The timing of the two stopping-related patterns is reminiscent of the times associated with reactive and proactive control, respectively (Braver, 2012; Braver et al., 2007; Chen et al., 2010; Chikazoe et al., 2009; Hanes et al., 1998; Ito et al., 2003; Majid et al., 2013; Stuphorn et al., 2000; Stuphorn et al., 2010; Stuphorn and Emeric, 2012). Moreover, by interleaving the tasks, we were able to show that it is largely the same neurons participating in both processes. Finally, we show that the patterns that differentiate value in the choice task can distinguish failed from successful stopping in the stop signal task. These results thus support the hypothesis that stopping and economic choice reflect common computations occurring in overlapping circuits.

These results provide evidence in favor of the hypothesis that the neural processes that regulate stopping relate to the ones that regulate economic choices. In foraging theory, decisions are generally framed as accept-reject (Blanchard and Hayden, 2015; Kacelnik et al., 2011; Stephens and Anderson, 2001; Stephens and Krebs, 1986). From this perspective, binary choices, the mainstay of behavioral economics and microeconomics, are better thought of as two somewhat independent accept-reject decisions (Hayden and Moreno-Bote, 2018; Kacelnik et al., 2011). Each accept-reject decision, in turn, functions like its classic foraging counterpart, that is, as a choice between pursuing and refraining from pursuit (Stephens and Krebs, 1986; Freidin et al., 2009; Kacelnik et al., 2011). In other words, what appears to be a binary choice may actually be a pair of countermanding stopping decisions (Hayden, 2018). Our results provide tentative neural evidence in support of this idea.

These results also invite a reconsideration of the role of the OFC. This region, especially Area 13, is sometimes cast as a specialist in economic functions (Padoa-Schioppa, 2011; Wallis, 2007). Our results challenge that narrow view and endorse a broader view that encompasses stopping as well. We doubt that the role of OFC is limited to these two functions. Its other functions likely include contingent (rule-based) decisions, working memory, switching, and conflict monitoring (Bryden and Roesch, 2015; Lara et al., 2009; Mansouri et al., 2014; Meyer and Bucci, 2016; Sleezer et al., 2016; Sleezer et al., 2017). All these functions, including stopping and economic choice, can arguably be placed within the larger category of executive functions. Like stopping, executive functions more broadly are generally associated with dorsal prefrontal structures (Miller, 2000; Miller and Cohen, 2001). Some of these functions may also be part of the repertoire of the OFC as well.

Stopping is often associated with dorsal prefrontal structures, such as FEF, and with subcortical structures, like SC (Hanes and Schall, 1995; Logan et al., 2015; Schall, 1991). Our work suggests that the role of OFC is qualitatively different than these regions. Specifically, we find evidence that single neurons were neither consistently associated with a higher or lower firing rate, nor were they associated with two discrete sets of neurons, as in frontal eye fields (FEF) and superior colliculus, SC (Hanes and Carpenter, 1999; Pouget et al., 2017; Stuphorn et al., 2000). Instead, our results show stopping correlates only when examining patterns found in ensembles of cells. This finding suggests that encoding of stop signals may be more abstract than the coding in more dorsal regions.

Overall, our results suggest one core function of OFC may be to generate an abstract regulatory signal to feed into a cascade of downstream structures that ultimately determine choice (Hunt and Hayden, 2017). In this way, it may be similar to other regions, especially cingulate cortex (Blanchard et al., 2015b; Hillman and Bilkey, 2010; Shenhav et al., 2013). In the context of economic choice, this signal will resemble a value signal; in other cases it will correlate with other relevant task variables. This view is consistent with the idea that choice and control processes both reflect a gradual transformation occurring in a distributed manner across brain regions, rather than a modular one (Balasubramani et al., 2018; Eisenreich et al., 2017; Hunt and Hayden, 2017) to study their neural codes. One benefit of view is that provides a basis for the observed role of OFC and adjacent structures in self-control (Kable and Glimcher, 2007; McClure et al., 2004).

The OFC is the major gateway by which sensory information enters into the prefrontal cortex and is a major source of visceral information as well (Öngür and Price, 2000). It may therefore occupy an early position in PFC processing hierarchies (Carmichael and Price, 1994; Fuster, 1988; Fuster, 2001; Rushworth et al., 2012; Rushworth et al., 2011). These facts raise the possibility that OFC serves as a first (or at least a relatively early) stage for computing preliminary executive signals that can affect-but not determine behavior (Cavada et al., 2000; Ebitz and Hayden, 2016; Öngür and Price, 2000; Wallis, 2007).

## METHODS

### Subjects

Two male rhesus macaques (Macaca mulatta, subject J, age 10, and subject T, age 5) served as subjects. All animal procedures were approved by the University Committee on Animal Resources at the University of Rochester and were designed and conducted in compliance with the Public Health Service’s Guide for the Care and Use of Animals.

### Recording site

A Cilux recording chamber (Crist Instruments) was placed over the area 13 of OFC (**Figure 1**). The targeted area expands along the coronal planes situated between 28.65 and 33.60 mm rostral to the interaural plane with varying depth. Position was verified by magnetic resonance imaging with the aid of a Brainsight system (Rogue Research Inc). Neuroimaging was performed at the Rochester Center for Brain Imaging, on a Siemens 3T MAGNETOM Trio Tim using 0.5 mm voxels. We confirmed recording locations by listening for characteristic sounds of white and grey matter during recording, which in all cases matched the loci indicated by the Brainsight system.

### Electrophysiological techniques

Single electrodes (Frederick Haer & Co., impedance range 0.8–4 MOhm) were lowered using a microdrive (NAN Instruments) until waveforms of between one and five neuron(s) were isolated. Individual action potentials were isolated on a Plexon system. Neurons were selected for study solely based on the quality of isolation; we never preselected based on task-related response properties.

### Eye tracking and reward delivery

Eye position was sampled at 1,000 Hz by an infrared eye-monitoring camera system (SR Research). Stimuli were controlled by a computer running MATLAB (Mathworks) with Psychtoolbox (Brainard and Vision, 1997) and Eyelink Toolbox (Cornelissen et al., 2002). A standard solenoid valve controlled the duration of water delivery. The relationship between solenoid open time and water volume was established and confirmed before, during, and after recording.

### Task

The stopping task is a measure of self-control that provides an alternative approach that avoids some of the limitations of intertemporal choice tasks (Hayden, 2016). The task followed standard stop signal paradigm (Hanes and Schall, 1995; Logan, 1994; Logan and Cowan, 1984). Subjects were placed in front of a computer monitor (1920x1080px) with black background. Following a brief (300 msec) central fixation on a white circle (radius 25px, **Figure 1**), the fixation spot disappeared on the appearance of eccentric saccade target (90px white square, 2.38 degrees, positioned at 288px in left or 1632px in right of screen, 50% chance). A go trial (67% of trials, randomly selected) was indicated by a go signal which is the peripheral target, whereas a stop trial (33% of trials, randomly selected) was indicated by an additional appearance of a stop signal — a central gray square (90px square, 2.38 degrees) delayed relative to the go signal presentation. Stop signal delays (SSD) in the task were set to stabilize at a delay causing approximately 50% successful stopping in average of all stop trials recorded till that moment of time in the day (SSD-50); SSDs were modulated through a staircase procedure with intervals of 16 msec. On go trials, subjects were rewarded for a saccade to the go signal and fixating on it for 200 msec; and on stop trials, subjects were rewarded for inhibiting their saccade to go signal and fixating at the stop signal for 400 msec. Water rewards were provided as feedback, and they were contingent on subject’s performance. Rewards were always 125 μl. The inter trial interval was 800 msec.

The *economic choice task* had a similar task framework to stop signal task, and they interleaved randomly in an interval of 1-3 trials. In go trials (random 67% of the total), a peripheral target called go offer (90px white square, 2.38 degrees, positioned at 288px in left or 1632px in right of the screen, 50% chance) was presented, and it was randomly associated with low (15μl), medium (125μl), or high (250μl) reward offers, as indicated by yellow, blue and magenta colored squares, respectively. In this task, the go trials were named forced choice trials, and the go offer was called offer 1. In stop trials (random 33% of the total)-called as choice trials, a center stop offer (offer 2, 90px square, 2.38 degrees) delayed with respect to the appearance of offer 1 was presented in addition. The offer 2 was also randomly associated with yellow, blue and magenta colors to indicate low, medium and high reward sizes. The offer 1 in stop trials was always in blue color to represent medium reward sized offer. This setup allowed the subject to make a choice through reward comparison in case of choice trials, and through a forced choice when only offer 1 was presented. All other parameters were the same as stop signal task.

### Behavioral analysis

Inhibition function related failed stoppings to stop signal delay (SSD). The delay from the presentation of go signal that caused 50% successful cancellation in stop signal task (SSD-50) was used for computing stop signal reaction time (SSRT). SSRT was usually computed through median and integration methods (Hanes and Schall, 1995; Logan, 1994; Logan and Cowan, 1984; Verbruggen and Logan, 2008). *Median* method computed median of go trials’ reaction time distribution and then subtracted SSD-50 from it to give SSRT. The *integration* method computed the point in go trials’ RT distribution whose area was half the whole and then subtracted SSD-50 from it to give SSRT. SSRT computed from both of the above methods gave nearly equal results, and they were averaged to obtain the final SSRT estimates reported for both subjects.

### Statistical methods

Separate PSTH matrices were constructed by aligning spike rasters to the presentation of the go signal and stop signal for every neuron. Firing rates were calculated in 1 msec bins but were generally analysed in longer epochs. Normalization procedure was carried out by subtracting the mean firing during inter-trial interval (ITI) time period (baseline) and then by zscoring each neuron’s data, and the normalized data is used for decoder analysis presented in the manuscript. Alternatively, decoding was also tested with just zscored data, and the results are presented in supplementary material. For display, PSTHs were smoothened using 200 msec running boxcars. Tests used in the study include two sample t-test for parameteric analysis, Wilcoxon rank test for non-parametric analysis, chi-square test for comparing decoder’s classification accuracy against baseline (50% classification accuracy), Pearson correlation method for correlation analysis. To compute population tuning, we picked neurons with significant (p < 0.05) differences between successful and failed stopping trials using Wilcoxon rank test.

### Decoding analyses

We chose a neural network based decoding technique because it could efficiently analyse population responses from frontal cortex that are highly multiplexed and nonlinear. To generate population activation states as input patterns for the decoding analysis, we first separated all trials of each neuron by trial conditions (successful and failed stopping trials). Then, we averaged the activity from randomly sampled 10 trials belonging to a condition, with replacement, to form activation state for a neuron in any particular time period. The averaged responses of all 96 neurons’ were pooled to generate one population activation state for a particular trial condition and for a specific time period. 100 unique activation state patterns were used for the network training. 75% of the data was used for training and the rest was used for testing the network. The procedure is similar to that carried out by other studies (e.g., Mante et al., 2013; Pouget et al., 2000; Rigotti et al., 2013; Wang and Hayden, 2017).

The network used to study the stopping patterns had a single hidden layer with 100 hidden nodes, and 2 output nodes each representing one target condition for classification. The number of input nodes equal to the total number of neurons used for analysis = 96 (from two subjects). The network weights were initialized to small random numbers between −0.01 and 0.01.

The following *back-propagation algorithm* was used for training the decoders (Haykin and Network, 2004; Rumelhart et al., 1986; Rumelhart et al., 1988; Werbos, 1974). In the below, the input nodes are denoted by subscript, *k*, hidden nodes by subscript, *j*, and output nodes by subscript, *i*. Output error, *e*, associated with the network’s response for the *p*’th input pattern was

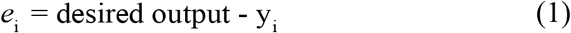

where *y*_i_ was the *i*’th output node response, and desired output was 1 / 0 if the *i*’th output node was associated with target trial condition for the corresponding input pattern (e.g., successful stopping, failed stopping). Total output error over all input patterns was computed by,

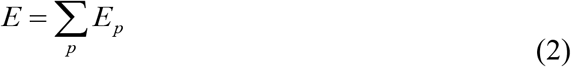

Network’s objective was to minimize the squared output error (eqn. 1) for the *p*’th pattern as denoted by eqn. (3).

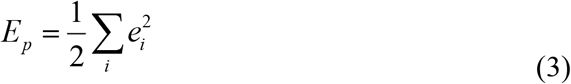

Response of any node was a hyperbolic tangent function (*g*) of slope = 5 of the total input (*h*_i_^s^) to it. The output node response, *y_i_*, as a function of its input was calculated as,

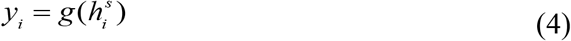

where, net input (*h*_i_^s^) to the output layer was,

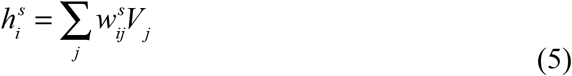

In the above, the weights, *w*_ij_, with superscript, *s*, indicate the second level of the network between hidden and output layer. *V*_j_ denoted the output of hidden layer, and it was represented as a function of net input to the hidden node (*h*_i_^f^) as follows,

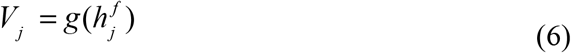

and

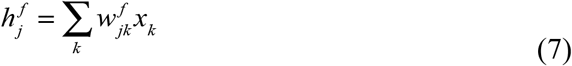

The superscript, *f*, in eqns. (6, 7) denote first level of the network between input and hidden layer, *w*_jk_ were their weights, and *x*_k_ was the input pattern to neural network. Weight updates were proportional to the negative change in error for the *p*’th pattern, *E_p_*, on change in weights. All updates happened trial by trial in the training phase. The update used at the second level was by eqn. (8), and that in the first level was by eqn. (10).

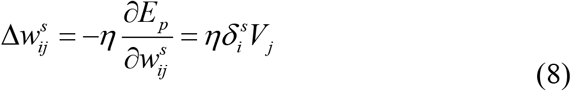

where,

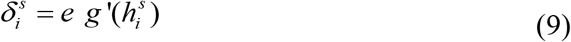

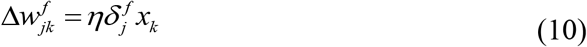

where,

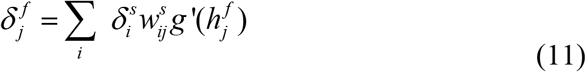

*ŋ* is the learning rate set to 0.001 for pre-go and post-stop signal decoder, and *g*’ denotes first order derivative of hyperbolic tangent function.

We had two different decoders trained on data from 1) pre-go signal, 2) post-stop signal time periods to show OFC’s active participation in stopping; the former worked on data aligned to presentation of go signal at time = 0, and the latter worked on data aligned to stop signal. For *pre-go decoder*, the training data was population activation states generated on averaging the signal from the fixation epoch spanning 300 msec before the presentation of go signal. For *post-stop decoder*, training data was generated on averaging the firing in the post-stop signal epoch. The entire network was run for *n* = 100 instances with different random weight initializations to obtain average output performance. Training procedure in all instances converged to classification accuracy of above 80%, and the converged weights at the end of training were used for testing of decoder. The testing data used were population activation states generated by averaging 100 msec boxcars that slides with step size of 10 msec (a total of 91 boxcars).

Similarities in the functioning and generalization of pre-go and post-stop decoders were analysed by comparing their converged weights, as well as by comparing their classification accuracy. For cross validation, the similarity index (r-max) was computed by cross correlating converged hidden layer weight vectors (with zero lag) of two decoders of interest. The index was averaged across *n* (=100) instances of networks with different weight initializations. The similarity index obtained from autocorrelating the weight vectors were used to statistically compare and cross-validate the results from cross correlation, and the results were significant using ttest (ttest, tstat = 210, p < 0.001). Comparisons between classification accuracies of the two decoders, at pre-go signal time period or post-stop signal time period, were done by using t-test on average performances of the two decoders computed from *n* instances (with different random weight initializations).

Cancellation time was defined by the size of test-boxcar window positioned at first instance of atleast four consecutive test-boxcars (100 msec window moving in intervals of 10 msec) in a row, whose performance was significantly higher than 50% using chi-square test (p < 0.05). The method avoids false positives that otherwise appear by 99% chance when considering just any one single significant instance of 91 total boxcars. With simulations using markov chains, we found that at least 4 consecutive significant windows were needed in a row for the claim of significance with p < 0.001; so the criteria to find at least 4 consecutive significant bins were used to find pre-go and post-stop decoder results as well as cancellation time. Average latency of cancellation signals to SSRT was found by subtracting SSRT of each subject from the mean cancellation time.

The decoders used for finding similarities between stopping and economic choices were similar to the above. The forced choice trials with offer 1 (accept) kind were chosen for analysis. The trials were divided into low and high value types, based on their reward magnitudes: the former type when the rewards were either low or medium, and the latter when the rewards were high, respectively. For this analysis, we also consider post-go signal decoder, similar to the post stop signal decoder except for its training on the post-go signal epoch from the presentation of go signal (in case of stop signal task) or offer 1 presentation (in case of economic choice task) till the reaction time. And, the feedback decoder was trained on the neural signals from the start to the end of feedback, for both the tasks. All decoders were tested using trials from economic choice task using boxcars of 100 msec length moving in intervals of 10 msec.

### Reward and stopping index

Reward index for every neuron was measured by linearly regressing the firing at outcome epoch (between reaction time and feedback) to the received reward sizes in *economic trials*. The stopping index was measured as the difference in normalized firing rates (FR) of successful and failed stopping trials divided by their norm.

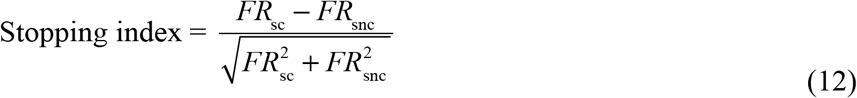

Cross validation tests were performed to support the idea that we had sufficient data to detect an effect had it been there, and to suggest that our results of lack of a significant correlation between stopping and reward indices were statistically meaningful. For the cross validation analysis, all trials within a neuron were randomly separated to two groups, A and B. Stopping and reward index were computed for those two groups of each neuron. We performed correlations between stopping indices of groups A and B, and between reward indices of A and B. A total of *n* (=100) random permutation instances were performed to generate different A and B sets. The test should ideally show high correlations between indices of A and B for any instance, and we indeed saw positive correlations between stopping-index_A_ and stopping-index_B_, and similarly for reward-index_A_ and reward-index_B_. We confirmed that the actual correlation coefficient between stopping and reward indices in OFC fell within bottom 1% of the coefficients computed for *n* instances of stopping-index_A_ and stopping-index_B_. The above was also confirmed for *n* coefficients for reward-index_A_ and reward-index_B_. The results showed no-significant correlations between stopping and reward indices with p ≤ 0.01.

## SUPPLEMENT

### Results-A

| Trial type |  | Reaction time distribution medians (s) |  | Encoding of value In feedback epoch |
| --- | --- | --- | --- | --- |
|  |  | *Subject J* | *Subject T* | *Both subjects* |
| Stopping task | Go trials | 0.43 ± 0.09 | 0.27 ± 0.12 | r = 0.27, p < 0.01<br>(unsigned: r = 0.35, p < 0.001) |
|  | Successful stopping trials | 0.39 ± 0.09 | 0.25 ± 0.08 |  |
|  | Failed stopping trials | 0.25 ± 0.20 | 0.15 ± 0.22 |  |
| Economic choice task | Forced choice trials | 0.41 ± 0.12 | 0.27 ± 0.13 | r = -1.5342, p = 0.0025 |
|  | Choose offer 2 trials | 0.46 ± 0.12 | 0.25 ± 0.10 |  |
|  | Choose offer 1 trials | 0.28 ± 0.19 | 0.20 ± 0.23 |  |

| Economic task Variables |  | *Subject J* | *Subject T* |
| --- | --- | --- | --- |
| Reaction times versus value of offers | Forced choice trials- offer 1 | Pearson correlation, r = 0.52, p < 0.001 | r = 0.43, p < 0.001 |
|  | Choice trials – offer 2 | r = 0.29, p = 0.003 | r = 0.27, p < 0.001 |
| Preference for offer 1 than offer 2 | Choice trials | 57.6%<br>(chi-square test, chi-square = 5.78, p = 0.016) | 56.25%<br>(chi-square stat = 3.38, p = 0.066) |
| Preference for offer 1 versus stop signal delay | Choice trials | slope of psychometric sigmoidal fit = 5.53 | Slope = 6.93 |
| Accuracy (choosing best offer) | Choice trials | 89.25% | 89.8% |
| Accuracy versus stop signal delay | Choice trials | slope of psychometric sigmoidal fit = -4.93 | Slope = -5.3 |
| Neuronal tuning for rewards | Offer 1 | 21.15% | 9.09% |
|  | Offer 2 | 11.5% | 9.09% |

Both subjects showed behavioral effects in the reaction times of successful stoppings as a function of previous trial conditions (**Figure 1**). Successful stopping trials were shorter when following a successful stopping trial (subject J: N = 328, subject T: N = 357) as opposed to following a failed stopping (subject J: N = 111, subject T: N = 60). The statistical t-test on data for subject J was 360 msec shorter, with t-stat = 11.33, p < 0.0001 and for subject T was: 290 msec shorter, t-stat = 11.88, p < 0.0001). Similarly, successful stopping trials were shorter when following a go trial (subject J: N = 833, subject T: N = 862) as opposed to following a failed stopping trial (subject J: 310 msec shorter, t-stat = 9.608, p < 0.0001 and subject T: 210 msec shorter, t-stat = 7.72, p < 0.0001). We found a choice of reject trials (choice of offer 2) were shorter when following another reject trial compared to an accept trial (subject J: 0.28 s, tstat = 8.47, p < 0.001, Figure K; subject T: 0.31 s, tstat = 9.49, p < 0.001, **Figure 1**). Consequent reject trials were also shorter than following an accept trial (subject J: 0.1 s, tstat = 5.96, p < 0.001, Figure K; subject T: 0.18, tstat = 4.97, p < 0.001). Accept trials were generally longer after following reject trials than following another accept trial (subject J: 0.05 s, tstat = 3.41, p < 0.001, Figure K; subject T: tstat = 2.06, p = 0.039).

The proportion of neurons that showed positive task-related tuning during successful stopping when regressed against baseline were 31.25%, while that showed negative tuning were 39.58%. Similarly, the proportion that showed positive task-related tuning during failed stopping activity when regressed against baseline were 34.38%, while that showed negative tuning were 34.38%. In economic choice task, the percent of neurons that showed positive feedback tuning when regressed against baseline were 45.83%, while that showed negative tuning were 51.04%. Overall, the tuning for success and failure of stopping significantly correlated with the feedback tuning of economic choice task (Pearson correlation, r = 0.62, p < 0.001).

## Results-B

### Population averages provide weak information about stopping

Analysis of single neurons did not provide strong evidence for a role for OFC in stopping. The percent of neurons that individually distinguish successful and failed stopping trials (regardless of sign) was 8.43% during the 100 msec post-stop signal time period, and was 10.50% during the 100 msec pre-go signal time period. (These epochs were selected before analysis in order to reduce the likelihood of p-hacking). These proportions were not significantly greater than chance in either of the two key epochs (chi-square stat = 1.22, p = 0.26 in the post-stop signal time period; chi-square stat = 1.8, p = 0.17 in the pre-go signal time period). This lack of a detectable effect does not imply that a correlation between stopping and unit activity in OFC does not exist; rather it suggests that if it does exist it is too weak to detect using conventional methods that focus on single neurons in a sample of the size we collected.

We next tested whether successful and failed stopping trials have a consistent sign of effect on firing rates. The percent of significantly positive cells (successful > failed) was 5.40%, and wasn’t significantly different from the percent of significantly negative (successful < failed) cells (3.03% chi-square test, chi-square stat = 0.52, p = 0.47) in the post-stop signal period. The difference in the sizes of the two cell classes was also not significant before the start of trial at the pre-go signal time period (significantly positive cells 7.55%, significantly negative cells 2.95%, chi square = 2.40, p = 0.12).

Next we looked at grand averages of populations of neurons. We observed no difference between successful and failed stopping trials either after the stop signal or before the beginning of trial. Specifically, during the post-stop signal time period, responses were slightly less for successful than failed stopping in subject J (average of 0.3 spikes/sec, p = 0.6); the opposite pattern was observed in subject T (average of 0.52 spikes/sec, p = 0.53). Neither effect was statistically significant. Thus, these results suggest that conventional population averages don’t reveal information about the pattern of stopping. Together these analyses indicate that, if stopping correlates exist in OFC, they are of a different form than they take in regions like FEF and SC.

**Figure.**
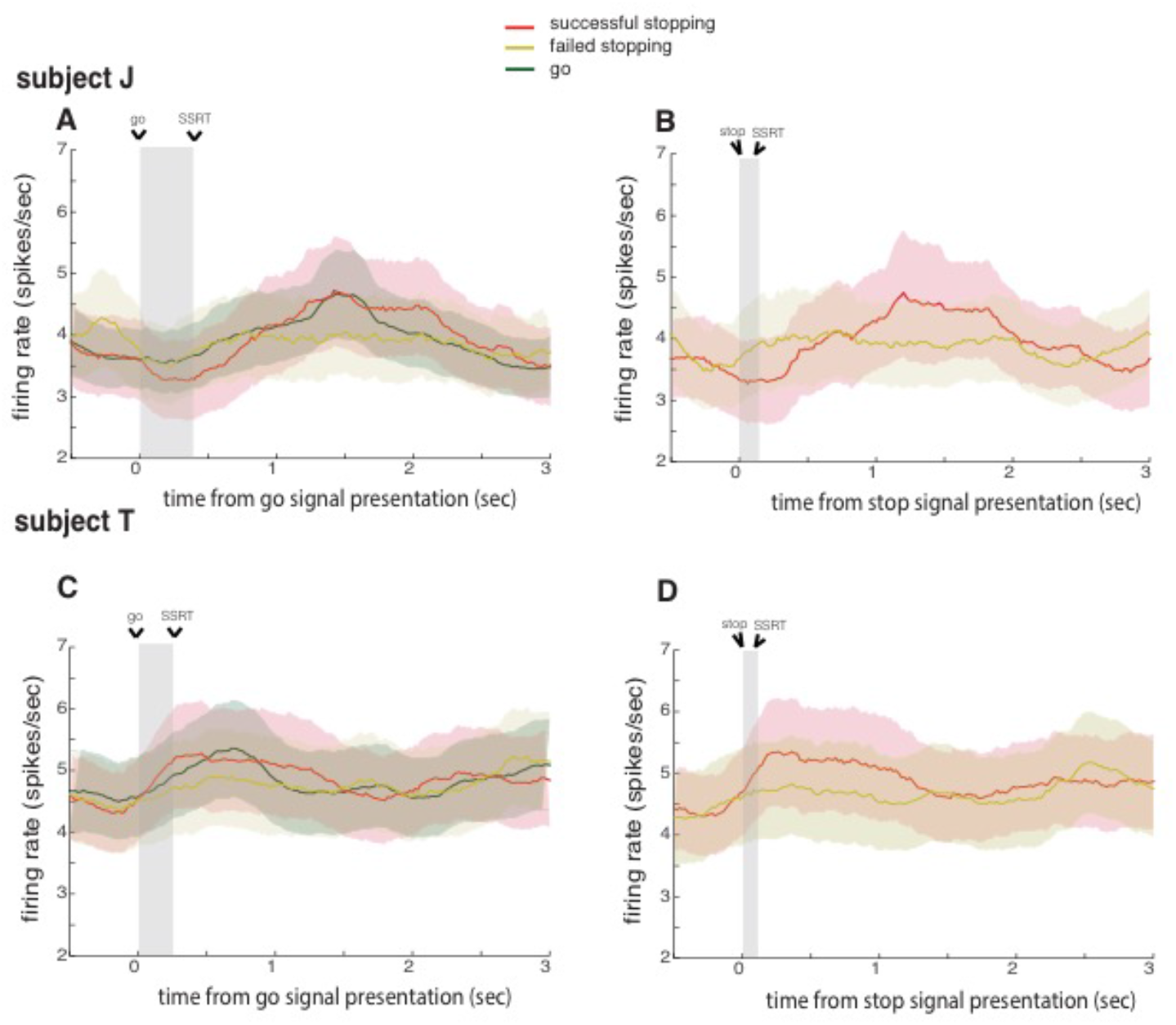
population averages provide weak information about stopping: Population activity for successful stopping and failed stopping with respect to **(A, C)** go signal presentation and **(B, D)** stop signal presentation, for subjects J and T. Time from start of the go (stop) signal to SSRT is shaded in panels A and C (B and D). Data for all SSDs are averaged to present successful and failed stopping trials. Error bars denote SEM. They don’t reveal significant information about the pattern of stopping.

## Results-C

**Figure.**
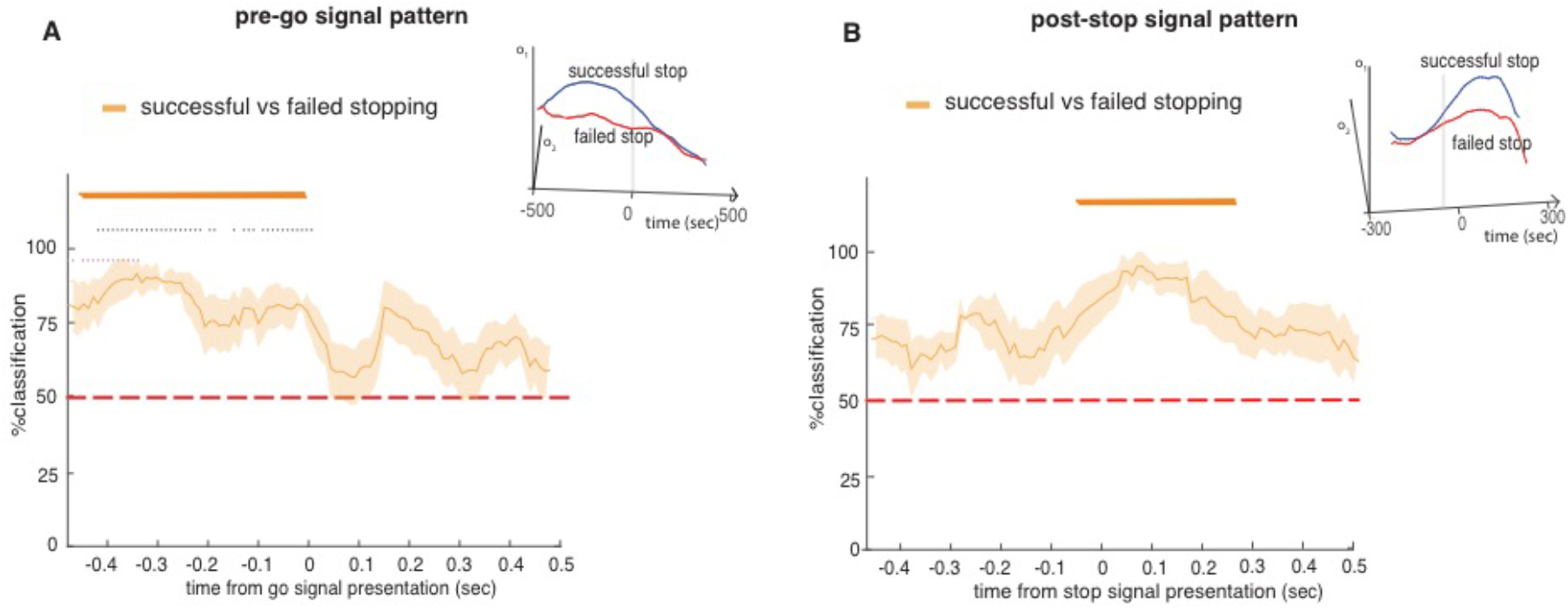
Post-stop and pre-go decoding for subjects J and T with zscored data (normalization): Both post stop signal and pre-go signal decoder was able to classify success of stopping significantly above chance (see **Methods** for specific use of chi-square tests to quantify significance) before SSRT and go signal presentation, respectively, and chi-square tests were used for finding their significance with p < 0.05, see **Methods**). Results suggest that successful differentiation of stopping codes can be obtained irrespective of the normalization methods used in the study (In the manuscript, Normalization procedure was carried out by subtracting the mean firing during inter-trial interval (ITI) time period (baseline) and then by zscoring each neuron**’**s data, and the normalized data is used for decoder analysis).

## Results-D

**Figure.**
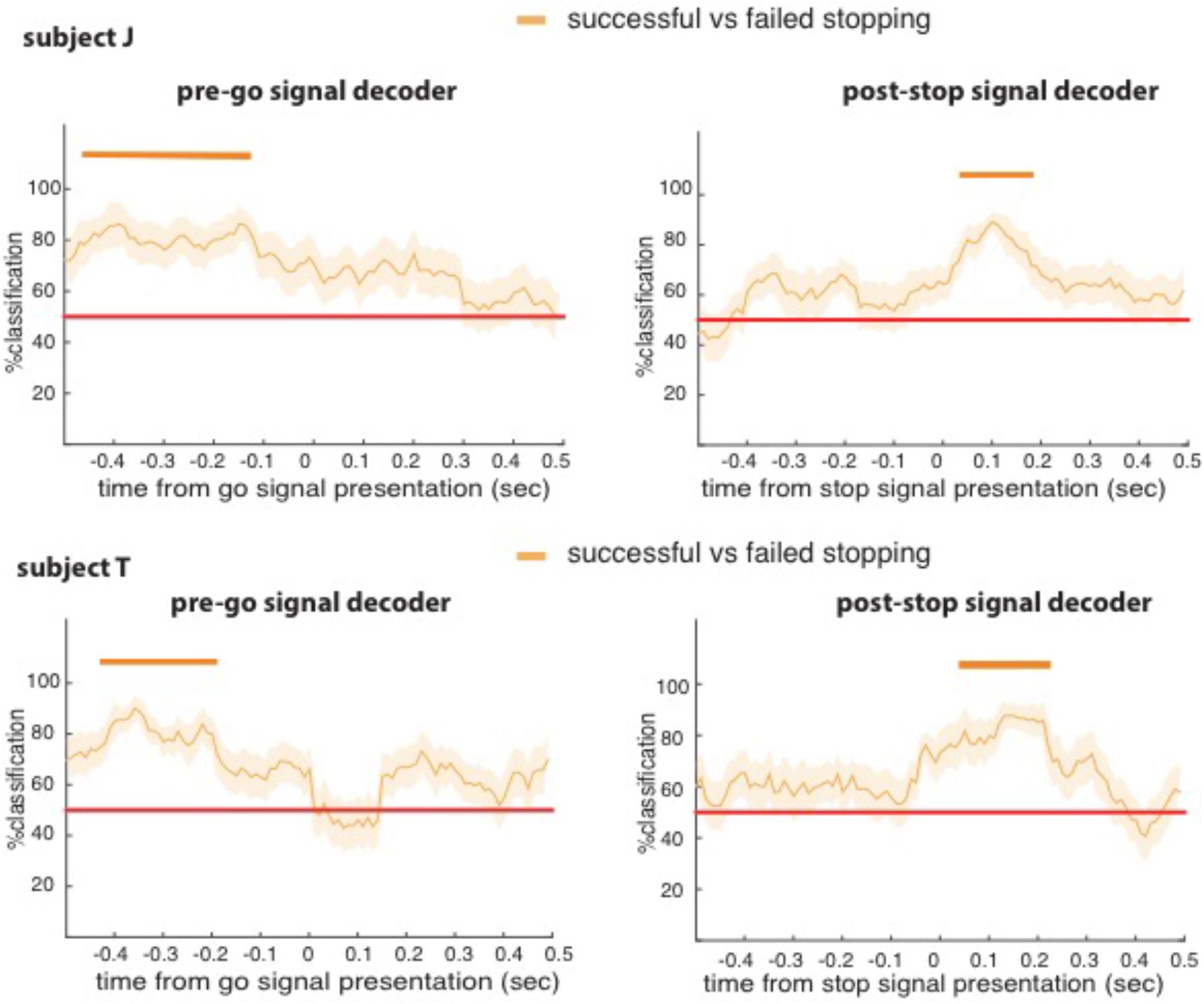
Post-stop and pre-go decoding for subjects J and T: The post stop signal decoder was able to classify success of stopping significantly above chance (see **Methods** for specific use of chi-square tests to quantify significance) in a time period ranging from 40 msec to 170 msec for subject J, and 40 to 220 msec for subject T, respectively after the stop signal (these times indicate the beginning of 100 msec boxcars, and chi-square tests were used for finding their significance with p < 0.05, see **Methods**). The first significant bin was therefore of window size 40 – 140 msec, that led to average cancellation time as 90 msec. It preceded the average stopping response by 50 msec in subject J, and by 30 msec in subject T, suggesting OFC**’**s responses may precede the stopping response. In Pre-go signal decoder, for subject J, high accuracy of decoding was found during the time periods 460 msec to 120 msec before the appearance of go signal. Likewise, it was 420 msec to 200 msec in subject T.

## Acknowledgements

This work was supported by an R01 (DA038615) to BYH. We thank Meghan C. Pesce for help with data collection.

## REFERENCES

Aron A.R. (2007). The neural basis of inhibition in cognitive control. The neuroscientist 13, 214–228.

Aron, A.R., and Poldrack, R.A. (2006). Cortical and subcortical contributions to stop signal response inhibition: role of the subthalamic nucleus. Journal of Neuroscience 26, 2424–2433.

Balasubramani, P.P., Moreno-Bote, R., and Hayden, B. (2018). Using a simple neural network to delineate some principles of distributed economic choice. Frontiers in Computational Neuroscience 12, 22.

Berkman, E., Hutcherson, C., Livingston, J.L., Kahn, L.E., and Inzlicht, M. (2016). Selfcontrol as value-based choice.

Blanchard, T.C., and Hayden, B.Y. (2015). Monkeys are more patient in a foraging task than in a standard intertemporal choice task. PloS one 10, e0117057.

Blanchard, T.C., Hayden, B.Y., and Bromberg-Martin, E.S. (2015a). Orbitofrontal cortex uses distinct codes for different choice attributes in decisions motivated by curiosity. Neuron 85, 602–614.

Blanchard, T.C., Strait, C.E., and Hayden, B.Y. (2015b). Ramping ensemble activity in dorsal anterior cingulate neurons during persistent commitment to a decision. Journal of neurophysiology 114, 2439–2449.

Brainard, D.H., and Vision, S. (1997). The psychophysics toolbox. Spatial vision 10, 433–436.

Braver T.S. (2012). The variable nature of cognitive control: a dual mechanisms framework. Trends in cognitive sciences 16, 106–113.

Braver, T.S., Gray, J.R., and Burgess, G.C. (2007). Explaining the many varieties of working memory variation: Dual mechanisms of cognitive control. Variation in working memory, 76–106.

Bryden, D.W., and Roesch, M.R. (2015). Executive control signals in orbitofrontal cortex during response inhibition. The Journal of Neuroscience 35, 3903–3914.

Carmichael, S., and Price, J. (1994). Architectonic subdivision of the orbital and medial prefrontal cortex in the macaque monkey. Journal of Comparative Neurology 346, 366–402.

Cavada, C., Compañy, T., Tejedor, J., Cruz-Rizzolo, R.J., and Reinoso-Suárez, F. (2000). The anatomical connections of the macaque monkey orbitofrontal cortex. A review. Cerebral Cortex 10, 220–242.

Chen, X., Scangos, K.W., and Stuphorn, V. (2010). Supplementary motor area exerts proactive and reactive control of arm movements. Journal of Neuroscience 30, 14657–14675.

Chikazoe, J., Jimura, K., Hirose, S., Yamashita, K.-i., Miyashita, Y., and Konishi, S. (2009). Preparation to inhibit a response complements response inhibition during performance of a stop-signal task. Journal of Neuroscience 29, 15870–15877.

Chudasama, Y., Kralik, J., and Murray, E. (2006). Rhesus monkeys with orbital prefrontal cortex lesions can learn to inhibit prepotent responses in the reversed reward contingency task. Cerebral Cortex 17, 1154–1159.

Cisek P. (2012). Making decisions through a distributed consensus. Current opinion in neurobiology 22, 927–936.

Cisek, P., and Kalaska, J.F. (2010). Neural mechanisms for interacting with a world full of action choices. Annual review of neuroscience 33, 269–298.

Cisek, P., and Pastor-Bernier, A. (2014). On the challenges and mechanisms of embodied decisions. Phil Trans R Soc B 369, 20130479.

Dias, R., Robbins, T., and Roberts, A. (1996). Dissociation in prefrontal cortex of affective and attentional shifts. Nature 380, 69–72.

Eagle, D.M., Baunez, C., Hutcheson, D.M., Lehmann, O., Shah, A.P., and Robbins, T.W. (2007). Stop-signal reaction-time task performance: role of prefrontal cortex and subthalamic nucleus. Cerebral cortex 18, 178–188.

Ebitz, R.B., and Hayden, B.Y. (2016). Dorsal anterior cingulate: a Rorschach test for cognitive neuroscience. Nature neuroscience 19, 1278.

Eisenreich, B.R., Akaishi, R., and Hayden, B.Y. (2017). Control without Controllers: Toward a Distributed Neuroscience of Executive Control. Journal of Cognitive Neuroscience.

Feierstein, C.E., Quirk, M.C., Uchida, N., Sosulski, D.L., and Mainen, Z.F. (2006). Representation of spatial goals in rat orbitofrontal cortex. Neuron 51, 495–507.

Freidin, E., Aw, J., and Kacelnik, A. (2009). Sequential and simultaneous choices: Testing the diet selection and sequential choice models. Behavioural processes 80, 218–223.

Fuster J.M. (1988). Prefrontal cortex. In Comparative neuroscience and neurobiology (Springer), pp. 107–109.

Fuster J.n.M. (2001). The prefrontal cortex—an update: time is of the essence. Neuron 30, 319–333.

Ghods-Sharifi, S., Haluk, D.M., and Floresco, S.B. (2008). Differential effects of inactivation of the orbitofrontal cortex on strategy set-shifting and reversal learning. Neurobiology of learning and memory 89, 567–573.

Grattan, L.E., and Glimcher, P.W. (2014). Absence of spatial tuning in the orbitofrontal cortex. PloS one 9, e112750.

Hampshire, A., and Sharp, D.J. (2015). Contrasting network and modular perspectives on inhibitory control. Trends in cognitive sciences 19, 445–452.

Hanes, D.P., and Carpenter, R. (1999). Countermanding saccades in humans. Vision research 39, 2777–2791.

Hanes, D.P., Patterson, W.F., and Schall, J.D. (1998). Role of frontal eye fields in countermanding saccades: visual, movement, and fixation activity. Journal of Neurophysiology 79, 817–834.

Hanes, D.P., and Schall, J.D. (1995). Countermanding saccades in macaque. Visual neuroscience 12, 929–937.

Hayden B.Y. (2018). Economic choice: the foraging perspective. Current Opinion in Behavioral Sciences 24, 1–6.

Hayden, B.Y., (2016). Time discounting and time preference in animals: A critical review. Psychon. Bull. Rev. 23, 39–53. https://doi.org/10.3758/s13423-015-0879-3

Hayden, B.Y., and Moreno-Bote, R. (2018). A neuronal theory of sequential economic choice. Brain and Neuroscience Advances 2, 2398212818766675.

Haykin, S., and Network, N. (2004). A comprehensive foundation. Neural Networks 2, 41.

Hillman, K.L., and Bilkey, D.K. (2010). Neurons in the rat anterior cingulate cortex dynamically encode cost-benefit in a spatial decision-making task. Journal of Neuroscience 30, 7705–7713.

Horn, N., Dolan, M., Elliott, R., Deakin, J., and Woodruff, P. (2003). Response inhibition and impulsivity: an fMRI study. Neuropsychologia 41, 1959–1966.

Hunt, L.T., and Hayden, B.Y. (2017). A distributed, hierarchical and recurrent framework for reward-based choice. Nat Rev Neurosci 18, 172–182.

Iacono, W.G., Malone, S.M., and McGue, M. (2008). Behavioral disinhibition and the development of early-onset addiction: common and specific influences. Annu Rev Clin Psychol 4, 325–348.

Inzlicht, M., Schmeichel, B.J., and Macrae, C.N. (2014). Why self-control seems (but may not be) limited. Trends in cognitive sciences 18, 127–133.

Ito, S., Stuphorn, V., Brown, J.W., and Schall, J.D. (2003). Performance monitoring by the anterior cingulate cortex during saccade countermanding. Science 302, 120122.

Kable, J.W., and Glimcher, P.W. (2007). The neural correlates of subjective value during intertemporal choice. Nature neuroscience 10, 1625.

Kacelnik, A., Vasconcelos, M., Monteiro, T., and Aw, J. (2011). Darwin’s “tug-of-war” vs. starlings’“horse-racing”: how adaptations for sequential encounters drive simultaneous choice. Behavioral Ecology and Sociobiology 65, 547–558.

Krajbich, I., Armel, C., and Rangel, A. (2010). Visual fixations and the computation and comparison of value in simple choice. Nature neuroscience 13, 1292.

Lara, A.H., Kennerley, S.W., and Wallis, J.D. (2009). Encoding of gustatory working memory by orbitofrontal neurons. Journal of Neuroscience 29, 765–774.

Lim, S.-L., O’Doherty, J.P., and Rangel, A. (2011). The decision value computations in the vmPFC and striatum use a relative value code that is guided by visual attention. Journal of Neuroscience 31, 13214–13223.

Logan G.D. (1994). On the ability to inhibit thought and action: A users’ guide to the stop signal paradigm.

Logan, G.D., and Cowan, W.B. (1984). On the ability to inhibit thought and action: A theory of an act of control. Psychological review 91, 295.

Logan, G.D., Yamaguchi, M., Schall, J.D., and Palmeri, T.J. (2015). Inhibitory control in mind and brain 2.0: blocked-input models of saccadic countermanding. Psychological review 122, 115.

Majid, D.A., Cai, W., Corey-Bloom, J., and Aron, A.R. (2013). Proactive selective response suppression is implemented via the basal ganglia. Journal of Neuroscience 33, 13259–13269.

Mansouri, F.A., Buckley, M.J., and Tanaka, K. (2014). The essential role of primate orbitofrontal cortex in conflict-induced executive control adjustment. Journal of Neuroscience 34, 11016–11031.

Mante, V., Sussillo, D., Shenoy, K.V., and Newsome, W.T. (2013). Context-dependent computation by recurrent dynamics in prefrontal cortex. nature 503, 78.

McClure, S.M., Laibson, D.I., Loewenstein, G., and Cohen, J.D. (2004). Separate neural systems value immediate and delayed monetary rewards. Science 306, 503–507.

Meyer, H.C., and Bucci, D.J. (2016). Imbalanced activity in the orbitofrontal cortex and nucleus accumbens impairs behavioral inhibition. Current biology 26, 2834–2839.

Miller E.K. (2000). The prefontral cortex and cognitive control. Nature reviews neuroscience 1, 59.

Miller, E.K., and Cohen, J.D. (2001). An integrative theory of prefrontal cortex function. Annual review of neuroscience 24, 167–202.

Mishkin M. (1964). Perseveration of central sets after frontal lesions in monkeys. The frontal granular cortex and behavior, 219–241.

Nestler, E.J., Barrot, M., DiLeone, R.J., Eisch, A.J., Gold, S.J., and Monteggia, L.M. (2002). Neurobiology of depression. Neuron 34, 13–25.

Ojeda, A., Murphy, R.A., and Kacelnik, A. (2018). Paradoxical choice in rats: subjective valuation and mechanism of choice. Behavioural processes.

Öngür, D., and Price, J. (2000). The organization of networks within the orbital and medial prefrontal cortex of rats, monkeys and humans. Cerebral cortex 10, 206–219.

Padoa-Schioppa C. (2011). Neurobiology of economic choice: a good-based model. Annual review of neuroscience 34, 333.

Padoa-Schioppa C. (2013). Neuronal origins of choice variability in economic decisions. Neuron 80, 1322–1336.

Padoa-Schioppa, C., and Assad, J.A. (2006). Neurons in the orbitofrontal cortex encode economic value. Nature 441, 223–226.

Pirrone, A., Azab, H., Hayden, B., Stafford, T., and Marshall, J.A. (2017). Evidence for the speed-value trade-off: human and monkey decision making is magnitude sensitive. Decision (in press).

Pouget, A., Dayan, P., and Zemel, R. (2000). Information processing with population codes. Nature Reviews Neuroscience 1, 125.

Pouget, P., Murthy, A., and Stuphorn, V. (2017). Cortical control and performance monitoring of interrupting and redirecting movements. Phil Trans R Soc B 372, 20160201.

Raghuraman, A.P., and Padoa-Schioppa, C. (2014). Integration of multiple determinants in the neuronal computation of economic values. Journal of Neuroscience 34, 11583–11603.

Rich, E., Stoll, F., and Rudebeck, P. (2017). Linking dynamic patterns of neural activity in orbitofrontal cortex with decision making. Current opinion in neurobiology 49, 24–32.

Rich, E.L., and Wallis, J.D. (2016). Decoding subjective decisions from orbitofrontal cortex. Nature neuroscience 19, 973.

Rigotti, M., Barak, O., Warden, M.R., Wang, X.-J., Daw, N.D., Miller, E.K., and Fusi, S. (2013). The importance of mixed selectivity in complex cognitive tasks. Nature 497, 585–590.

Roberts, A., and Wallis, J. (2000). Inhibitory control and affective processing in the prefrontal cortex: neuropsychological studies in the common marmoset. Cerebral Cortex 10, 252–262.

Roesch, M.R., Taylor, A.R., and Schoenbaum, G. (2006). Encoding of time-discounted rewards in orbitofrontal cortex is independent of value representation. Neuron 51, 509–520.

Rudebeck, P.H., and Murray, E.A. (2014). The orbitofrontal oracle: cortical mechanisms for the prediction and evaluation of specific behavioral outcomes. Neuron 84, 1143–1156.

Rumelhart, D., McClelland, J., and Williams, R. (1986). Parallel recognition in modern computers. processing: Explorations in the microstructure of cognition 1.

Rumelhart, D.E., McClelland, J.L., and Group, P.R. (1988). Parallel distributed processing, Vol 1 (IEEE).

Rushworth, M.F., Kolling, N., Sallet, J., and Mars, R.B. (2012). Valuation and decisionmaking in frontal cortex: one or many serial or parallel systems? Current opinion in neurobiology 22, 946–955.

Rushworth, M.F., Noonan, M.P., Boorman, E.D., Walton, M.E., and Behrens, T.E. (2011). Frontal cortex and reward-guided learning and decision-making. Neuron 70, 1054–1069).

Sakagami, M., and Pan, X. (2007). Functional role of the ventrolateral prefrontal cortex in decision making. Current opinion in neurobiology 17, 228–233.

Schall J.D. (1991). Neuronal activity related to visually guided saccades in the frontal eye fields of rhesus monkeys: comparison with supplementary eye fields. Journal of neurophysiology 66, 559–579.

Schall J.D. (2001). Neural basis of deciding, choosing and acting. Nature Reviews Neuroscience 2, 33–42.

Schall, J.D., Stuphorn, V., and Brown, J.W. (2002). Monitoring and control of action by the frontal lobes. Neuron 36, 309–322.

Schoenbaum, G., Roesch, M.R., Stalnaker, T.A., and Takahashi, Y.K. (2009). A new perspective on the role of the orbitofrontal cortex in adaptive behaviour. Nature Reviews Neuroscience 10, 885–892.

Schoenbaum, G., Setlow, B., Nugent, S.L., Saddoris, M.P., and Gallagher, M. (2003). Lesions of orbitofrontal cortex and basolateral amygdala complex disrupt acquisition of odor-guided discriminations and reversals. Learning & Memory 10, 129–140.

Schoenbaum, G., Takahashi, Y., Liu, T.L., and McDannald, M.A. (2011). Does the orbitofrontal cortex signal value? Annals of the New York Academy of Sciences 1239, 87–99.

Schultz W. (2000). Multiple reward signals in the brain. Nature reviews Neuroscience 1, 199.

Shapiro, M.S., Siller, S., and Kacelnik, A. (2008). Simultaneous and sequential choice as a function of reward delay and magnitude: normative, descriptive and process-based models tested in the European starling (Sturnus vulgaris). Journal of Experimental Psychology: Animal Behavior Processes 34, 75.

Shenhav A. (2017). The perils of losing control: Why self-control is not just another value-based decision. Psychological Inquiry

Shenhav, A., Botvinick, M.M., and Cohen, J.D. (2013). The expected value of control: an integrative theory of anterior cingulate cortex function. Neuron 79, 217–240.

Sleezer, B.J., Castagno, M.D., and Hayden, B.Y. (2016). Rule encoding in orbitofrontal cortex and striatum guides selection. Journal of Neuroscience 36, 11223–11237.

Sleezer, B.J., LoConte, G.A., Castagno, M.D., and Hayden, B.Y. (2017). Neuronal responses support a role for orbitofrontal cortex in cognitive set reconfiguration. European Journal of Neuroscience 45, 940–951.

Stalnaker, T.A., Cooch, N.K., and Schoenbaum, G. (2015). What the orbitofrontal cortex does not do. Nature Neuroscience 18, 620–627.

Stephens, D.W., and Anderson, D. (2001). The adaptive value of preference for immediacy: when shortsighted rules have farsighted consequences. Behavioral Ecology 12, 330–339.

Stephens, D.W., and Krebs, J.R. (1986). Foraging theory (Princeton University Press).

Strait, C.E., Blanchard, T.C., and Hayden, B.Y. (2014). Reward value comparison via mutual inhibition in ventromedial prefrontal cortex. Neuron 82, 1357–1366.

Strait, C.E., Sleezer, B.J., and Hayden, B.Y. (2015). Signatures of value comparison in ventral striatum neurons. PLoS Biol 13, e1002173.

Strait CE, Sleezer BJ, Blanchard TC, Azab H, Castagno MD, Hayden BY. (2016). Neuronal selectivity for spatial positions of offers and choices in five reward regions. Journal of neurophysiology 115(3):1098–111.

Stuphorn, V., Bauswein, E., and Hoffmann, K.-P. (2000). Neurons in the primate superior colliculus coding for arm movements in gaze-related coordinates. Journal of Neurophysiology 83, 1283–1299.

Stuphorn, V., Brown, J.W., and Schall, J.D. (2010). Role of supplementary eye field in saccade initiation: executive, not direct, control. Journal of neurophysiology 103, 801–816.

Stuphorn, V., and Emeric, E.E. (2012). Proactive and reactive control by the medial frontal cortex. Frontiers in Neuroengineering 5.

Sugrue, L.P., Corrado, G.S., and Newsome, W.T. (2005). Choosing the greater of two goods: neural currencies for valuation and decision making. Nature Reviews Neuroscience 6, 363.

Vasconcelos, M., Monteiro, T., Aw, J., and Kacelnik, A. (2010). Choice in multialternative environments: a trial-by-trial implementation of the sequential choice model. Behavioural processes 84, 435–439.

Verbruggen, F., and Logan, G.D. (2008). Response inhibition in the stop-signal paradigm. Trends in cognitive sciences 12, 418–424.

Volkow, N.D., Wang, G.-J., Fowler, J.S., Tomasi, D., and Telang, F. (2011). Addiction: beyond dopamine reward circuitry. Proceedings of the National Academy of Sciences 108, 15037–15042.

Wallis J.D. (2007). Orbitofrontal cortex and its contribution to decision-making. Annu Rev Neurosci 30, 31–56.

Werbos P. (1974). Beyond regression: New tools for prediction and analysis in the behavioral sciences.

Wang, M.Z., and Hayden, B.Y. (2017). Reactivation of associative structure specific outcome responses during prospective evaluation in reward-based choices. Nature Communications 8, ncomms15821.

Wilson, R.C., Takahashi, Y.K., Schoenbaum, G., and Niv, Y. (2014). Orbitofrontal cortex as a cognitive map of task space. Neuron 81, 267–279.

Xie, Y., Nie, C., and Yang, T. (2018). Covert shift of attention modulates the value encoding in the orbitofrontal cortex. eLife 7, e31507

Yoo, S.B.M., Sleezer, B.J., and Hayden, B.Y. (2018). Robust Encoding of Spatial Information in Orbitofrontal Cortex and Striatum. Journal of cognitive neuroscience, 1–16.

